# Suppression and modulation mechanism underlying the flavonoid-induced inhibition of fibrillation of α-synuclein

**DOI:** 10.1101/2025.08.21.671593

**Authors:** Geetika Verma, Rajiv Bhat

## Abstract

The accumulation of α-synuclein (αSyn) aggregates is a hallmark of synucleinopathies, including Parkinson’s disease (PD). However, the progression from harmless monomeric αSyn to misfolded oligomers and fibrillar species remains largely unclear. In this study, we examined the effect of Cyanidin, a naturally occurring neuroprotective compound, on the aggregation properties of α-syn using a combination of various biophysical tools. Thioflavin T fluorescence measurements revealed the inhibition of α-syn fibrillation in the presence of sub-stoichiometric concentration of Cyanidin, indicating its effect on α-syn nucleation. Ultracentrifugation and size exclusion chromatographic analyses demonstrated that increasing concentrations of Cyanidin reduced the conversion of monomers to aggregated forms of α-syn. Analysis of the aggregation reaction of α-syn based on monomer concentration suggested that Cyanidin reduces the *in-vitro* conversion of monomers to amyloid nuclei at sub-stoichiometric micromolar concentrations. Cyanidin was observed to bind weakly with the unstructured, monomeric α-syn, limiting the nucleation step and modulating the pathway to form conformationally restrained, SDS-resistant α-helical higher-order oligomers (∼670 kDa), which are less-hydrophobic. AFM and TEM images show that Cyanidin also possesses strong fibril disaggregation activity. Furthermore, seeding studies reveal that the higher-order oligomers, which are otherwise stable, destabilise upon sonication and attain seeding capability. Steady-state fluorescence spectroscopy and isothermal titration calorimetry (ITC) reveal weak interactions between Cyanidin and αSyn, with a dissociation constant in the mM range. Interestingly, Cyanidin-generated oligomers increase the viability of neuroblastoma (SH-SY5Y) cells, thereby protecting the neuronal cells from degeneration. Our study suggests that stabilization of structured oligomers by small molecule modulators like Cyanidin provides a viable strategy to interfere with αSyn fibrillization.

## Introduction

Parkinson’s disease (PD), a progressive neurodegenerative disorder affecting over 6.1 million people globally is characterized by cardinal motor symptoms including tremor, postural instability, bradykinesia, and muscular rigidity ^1^. A plethora of evidences suggest that the presence of fibrillar Lewy body deposits within the dopaminergic neurons of the substantia nigra pars compacta of the mid-brain contributes to neuronal cell death and disease progression in PD. Later, an unstructured and intrinsically disordered protein alpha-Synuclein (α-Syn) was identified as a major component of these inclusions^2^. α-Syn, a 140 amino acid with a molecular mass of ∼14.4 kDa, is a natively unfolded protein that is mainly expressed in the pre-synaptic terminal and is known to be involved in neuronal plasticity, synaptic vesicle trafficking, and neurotransmitter release. In vitro, α-Syn has been observed to follow the nucleation-conversion-polymerization model of aggregation^3^. The kinetics of this aggregation pathway initiates with the primary nucleation phase, where the native monomeric α-Syn protein forms ‘pre-nuclei’, which then convert into oligomer seeds (nuclei), that eventually elongate via monomer addition, to give rise to fibrils^4^.

According to current hypothesis, the initial oligomeric intermediates of fibrillar structures that are common to various disease-related neurodegenerative proteins including α-Syn, are the primary toxic species which upon maturing into fibrils are otherwise less toxic. The fibrils further fragment into oligomers which act as ‘seeds’ or ‘secondary nuclei’ for de novo fibrillation on existing fibril surfaces, leading to a loop of fibril amplification^5, 6^. Owing to their structural heterogeneity, a comprehensive understanding of the structural basis of the pathogenicity of these oligomeric conformations is not widely known, mainly because of their transient nature, heterogeneity and apparent low concentration at any particular time-point of the fibrillation pathway, thereby considerably hampering their investigation.

The aberrant assembly of oligomeric intermediates could be toxic or non-toxic in nature. Several findings suggest that structural flexibility and hydrophobic exposure are primary determinants of the ability of oligomeric assemblies to cause cellular dysfunction and eventually lead to neurodegeneration ^7^. Rising evidence of toxicity associated with the oligomeric species formed during amyloidogenesis has shifted the focus from inhibiting amyloid plaque formation to developing strategies that could modulate the species to become less toxic. Several studies show that various small molecules, such as flavones (e.g., baicalein) and polyphenols (e.g., epigallocatechin gallate), drift α-Syn towards the formation of non-toxic oligomeric species^8^. In order to understand the effects of α-Syn oligomers formed during its course of fibrillation and prevent the onset of underlying diseases, it is important to gain an in-depth understanding of the effects of such modulators on the oligomerization and fibrillation pathway of α-Syn. One of the biggest challenges in understanding oligomers is obtaining them in a stable form. This difficulty arises from the nature of these oligomers, which are extremely labile and short-lived ^9^. Once formed and reaching a threshold concentration, they are rapidly converted into mature fibrils. Current studies focus on studying the oligomeric species formed during the aggregation process in the presence of the flavone Cyanidin.

In our study, we delve into the modulating effects of Cyanidin, a known anti-amyloidogenic flavonoid, on the oligomerization and fibrillation propensity of α-Syn. The results shed light on the mechanisms underlying fibrillation inhibition by Cyanidin. Our prior investigations involving three anthocyanidins–Delphinidin, Peonidin and Cyanidin– revealed differences in their anti-amyloidogenic properties and showed variations in the morphology of potentially non-toxic species formed in the presence of Peonidin and Cyanidin. While we have elucidated the mechanism of α-syn fibrillation leading to amorphous aggregates in the presence of Peonidin (Ref.), the mechanism of the modulatory effect of Cyanidin has not been explored.

With this background, the current work presents findings on the structure, size, stability, and potential cellular impact of *in vitro* α-Syn higher-order oligomer-like structures formed in the presence of Cyanidin using a range of biophysical techniques. In the presence of Cyanidin, the oligomers formed during α-Syn fibrillation exhibit unique characteristics compared with those formed in its absence. Specifically, Cyanidin, at a sub-stoichiometric concentration as low as 40 μM, considerably slows the formation of fibril nuclei, thereby inhibiting fibril polymerization by modulating the ongoing fibrillation pathway. It promotes the formation of SDS-resistant higher-order α-Syn oligomers (approximately 15-mer) that adopt a conformationally constrained α-helical structure during fibrillation. Notably, this effect arises from weak, non-covalent interactions (with a dissociation constant, K_d_, in the millimolar range) between Cyanidin and monomeric α-Syn. Docking analysis indicates that Cyanidin predominantly binds to the N-terminus and the hydrophobic NAC domain of α-Syn. TEM images reveal intriguing associations of protein molecules to form droplet-shaped structures when α-Syn interacts with Cyanidin. Interestingly, these Cyanidin-induced α-Syn droplets are SDS-stable, non-hydrophobic, and non-toxic in nature. From a thermodynamic standpoint, the mesh-like assembly induced by Cyanidin slows down the conformational evolution along the pathway leading to the fibrillar amyloid state. In summary, Cyanidin stabilizes these oligomer-like droplet-shaped higher-order structures of α-Syn. These findings provide valuable insights into the interplay between the flavonoid Cyanidin and α-Syn, with potential implications for PD.

## Results

### Cyanidin inhibits fibrillation and redirects α-Syn towards non-fibrillar aggregates

In previous studies, Cyanidin has been identified as one of the flavonoid molecules that inhibits fibrillation of α-synuclein. However, the mechanism behind this effect remains unknown. The inhibitory effect of Cyanidin on the aggregation kinetics of α-Syn was analysed by thioflavin-T (ThT) binding assay (Figure 1A). ThT is a benzothiazole dye that fluoresces strongly upon binding with the cross-β-sheet structures, which are a signature of amyloid fibrils. Th-T fluorescence emission was monitored during the aggregation kinetics of α-Syn (70 μM) in the absence and presence of different concentrations of Cyanidin (5 μM to 200 μM). Consistent with our previous work, a concentration-dependent decrease in ThT fluorescence emission was observed, where concentrations of 30 μM and above showed a complete inhibition of fibrillation. The addition of as little as ∼0.4 equivalents of Cyanidin (i.e., 30 μM) delayed the aggregation with a significantly longer *t*_lag_ of 10 hours, while ∼0.7 equivalents of Cyanidin (i.e., 50 μM) completely abolished α-syn fibril formation (Figure 1A, B). The decrease in the apparent fibrillation rate (*k*_app_) of α-Syn by the addition of Cyanidin is shown in Figure 1B. To further establish that the decrease in ThT fluorescence in the presence of Cyanidin is due to fibril inhibition and not due to interference between ThT and Cyanidin, the negative control fluorescence was also monitored. No significant change in the ThT fluorescence intensity was observed in the presence of Cyanidin, suggesting that no interaction occurs between Cyanidin and ThT (Figure S1).

**Figure 1:**
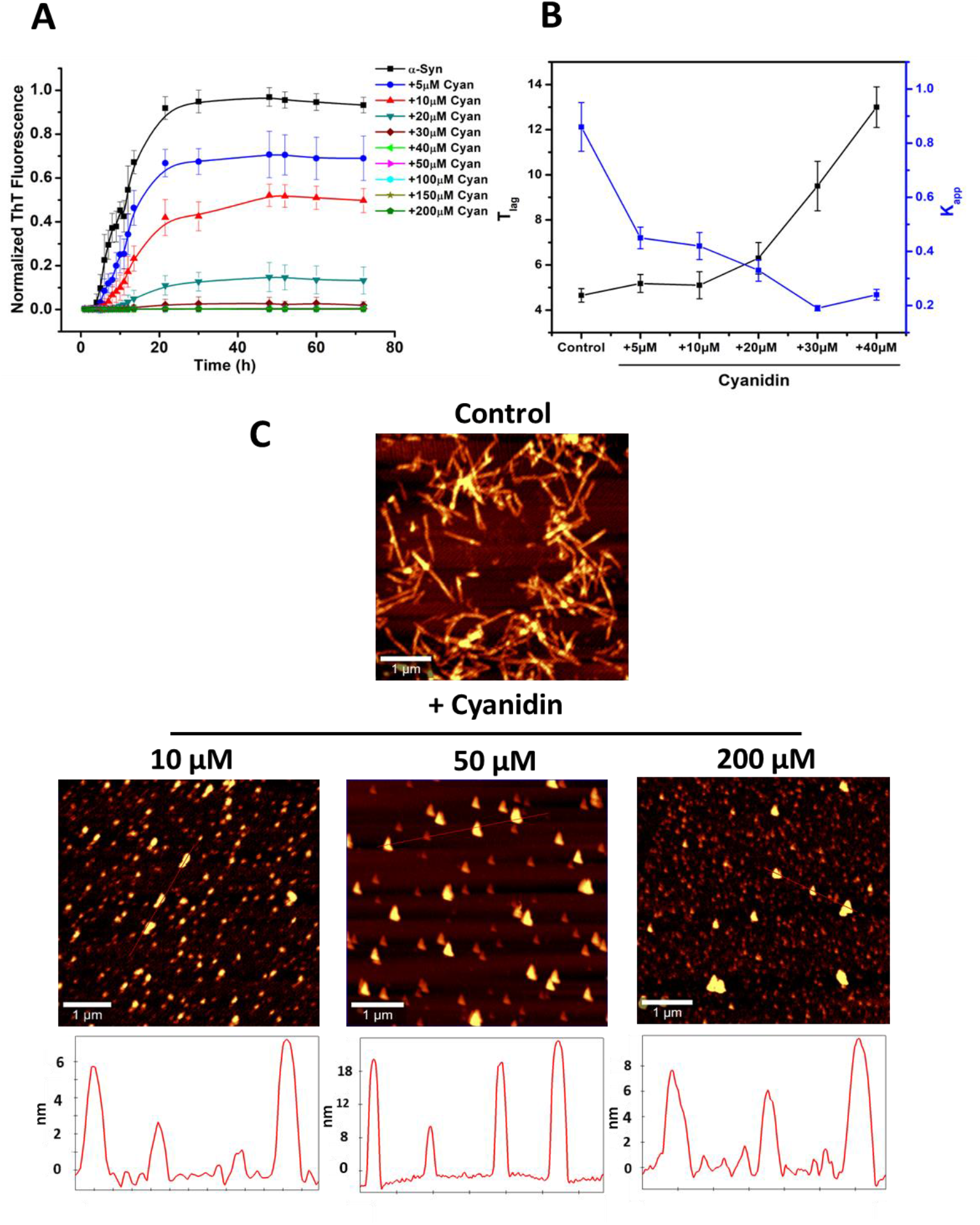
Cyanidin inhibits α-Syn fibrillation in a concentration-dependent manner. **(A)** ThT Fibrillation kinetics of α-Syn (70 μM) studied in the presence of increasing concentrations of Cyanidin (5 – 200 μM) for 72 hours at 37^°^C under continuous shaking conditions at 200 rpm in an incubator shaker. **(B)** The graph represents the change in lag time (T_lag_) and apparent rate constant of fibrillation (k_app_) with increasing concentrations of Cyanidin. The error bars represent the standard deviation from three independent aggregation reactions, ±SD (n = 3). **(C)** Atomic force microscopy images showing significant appearance of oligomeric α-Syn species in the presence of different concentrations of Cyanidin.

Interestingly, Cyanidin completely suppresses α-Syn fibrillation at sub-stoichiometric concentrations, with a ratio of less than 1:1 (protein:inhibitor) bringing about complete inhibition (Figure 1A). The strong inhibition of α-Syn fibrillation by a much lower concentration of Cyanidin indicates that it targets the nucleation phase of the pathway, interacting with the nucleus or seeds present in low concentrations, similar to how small molecules inhibit Aβ fibrillation^10^. The morphologies of α-syn aggregates formed in the presence of different concentrations of Cyanidin were visualised by atomic force microscopy (AFM). Figure 1C shows AFM images of α-Syn fibrils formed both in the absence and presence of Cyanidin, demonstrating a significant suppression of fibril formation in the presence of Cyanidin. α-Syn (70 μM) in the absence of Cyanidin formed mature fibrils with an average height of ∼9 nm, while no fibrils were observed in the presence of Cyanidin. However, it has been observed that α-syn aggregates formed had different morphologies after incubation for 72h with three different concentrations of Cyanidin – 10, 50, and 200 μM, with an average height ranging from ∼8 to 20 nm. This suggests that sub-stoichiometric concentrations of Cyanidin have a strong propensity to structurally transform α-syn into conformations other than fibrils. Interestingly, at 50 μM concentration, Cyanidin generates oligomer-like higher-order structures which appear to have droplet shapes unlike the fibrillar structure formed in the absence of Cyanidin during the aggregation of α-syn^11^ (Fig. 2C). The on-pathway α-syn oligomers formed during the course of fibrillation pathway (in the absence of Cyanidin) (Figure S2) were compared with the Cyanidin-induced droplet shaped oligomers (Figure 1C), and AFM revealed that the two oligomeric species could be categorized into two groups based on their diameters: 4-8 nm and 18-20 nm, respectively (Figure S2 and Figure 1C, respectively). These dimensions of oligomers formed in the absence of Cyanidin are consistent with those previously identified for on-pathway oligomers using AFM^12^.

**Figure 2:**
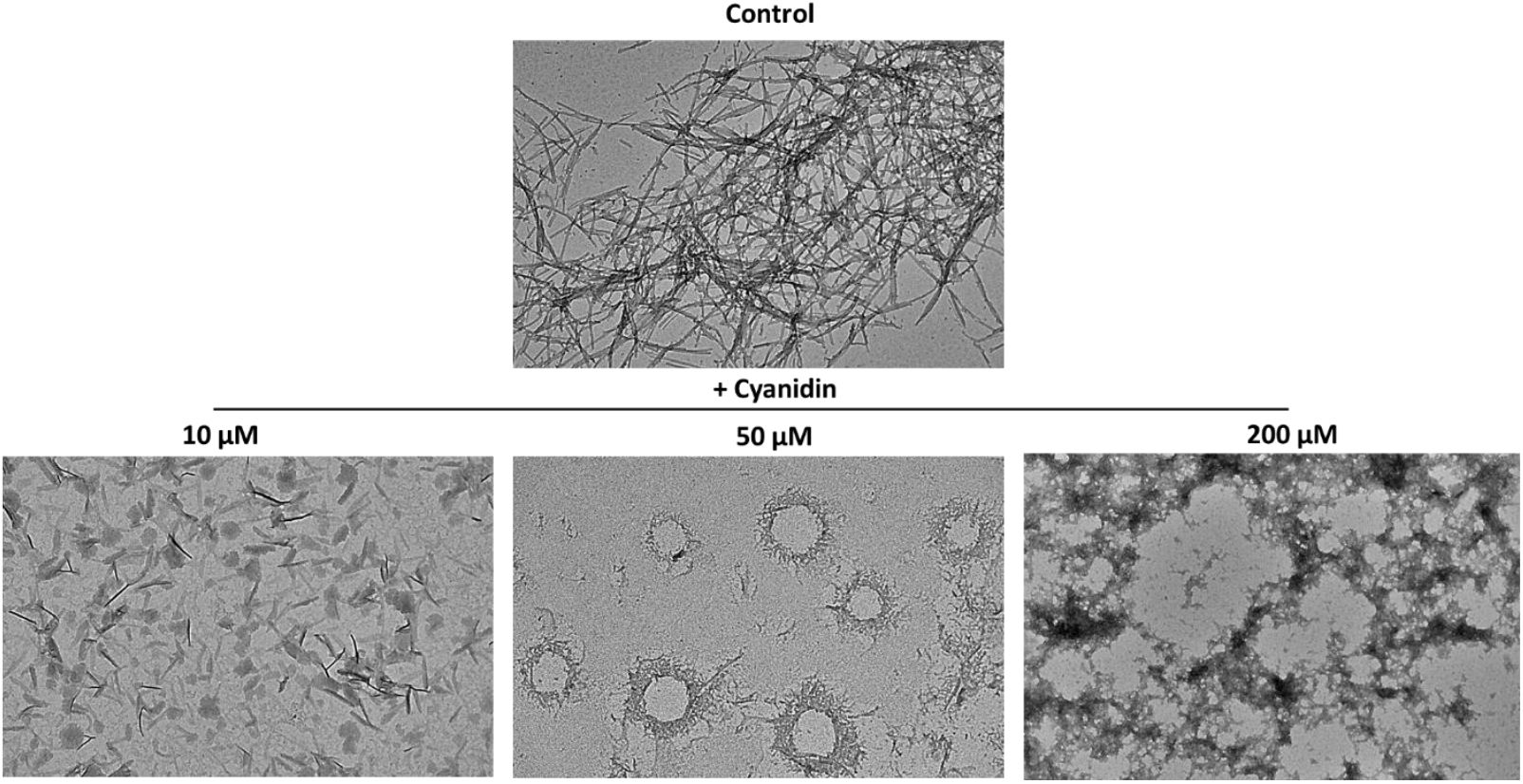
Transmission electron microscopy (TEM) images showing varying patterns of aggregation of α-Syn. – α-syn fibrils (Control) and α-Syn aggregates formed in the presence of different concentrations (10, 50, and 200 µM) of Cyanidin.

In addition, we employed transmission electron microscopy (TEM) to complement the AFM results. The TEM image of the control sample (α-syn without Cyanidin) reveals the presence of dense fibrils (Figure 2). In contrast, when Cyanidin is introduced at different concentrations, shorter fibrils become evident at 10 µM Cyanidin, hollow-shaped clusters of higher-order aggregates are prominent at 50 µM Cyanidin, and amorphous aggregates are observed at 200 µM Cyanidin. The hollow-shaped α-syn oligomer-like aggregates formed in the presence of 50 µM Cyanidin were of different dimensions, as shown in Figure S2. Nonetheless, different Cyanidin concentrations are observed to strongly affect the size and shape distribution of α-Syn aggregates.

### Cyanidin disaggregates mature α-Syn fibrils and attenuates fibril elongation

Disaggregation of amyloid fibrils can be investigated either by indirectly monitoring the reduction in ThT intensity or directly observing via AFM or TEM. ThT fluorescence was, hence, monitored upon addition of 50 μM Cyanidin at different time-intervals (0, 3, 6, 12, 24 and 48h) of the fibrillation pathway. The ThT fluorescence was found to drop instantly after the addition of Cyanidin at different time-points (Figure 3A), indicating the replacement of ThT molecules by Cyanidin or the shielding of ThT emission by Cyanidin molecules. Therefore, it is difficult to accurately evaluate the disassembly activity of Cyanidin toward α-Syn amyloid fibrils using only ThT assay. After the addition of Cyanidin at different time points, the overall size estimation of the α-Syn species was monitored using static light scattering (Figure 3B). The results indicated that Cyanidin was effective in inhibiting α-Syn fibrillation when added up to the initial log phase (i.e., 6 hours) of the fibrillation pathway. However, when added at later stages, it led to greater scattering, indicating the presence of structures similar to fibrils that did not bind to ThT. Morphological observations using AFM and TEM directly demonstrated the disaggregation activity of Cyanidin against α-Syn fibrils. Both AFM and TEM images revealed the presence of shorter and disintegrated fibril fragments, indicating the disaggregation of protofibrils and pre-formed α-Syn fibrils after the addition of Cyanidin during the log and saturation phases (12 h, 24 h, and 48 h, respectively) of the α-Syn fibrillation pathway (Figure 3C and D, respectively). Examples in the literature have earlier shown that small naturally occurring compounds can disaggregate amyloid fibrils. Simulation studies have also shown that these compounds disrupt the salt bridge, causing instability in the hydrogen bonds within the amino acid backbone, which supports the fibrils, ultimately leading to their remodelling or disaggregation ^13,14^.

**Figure 3:**
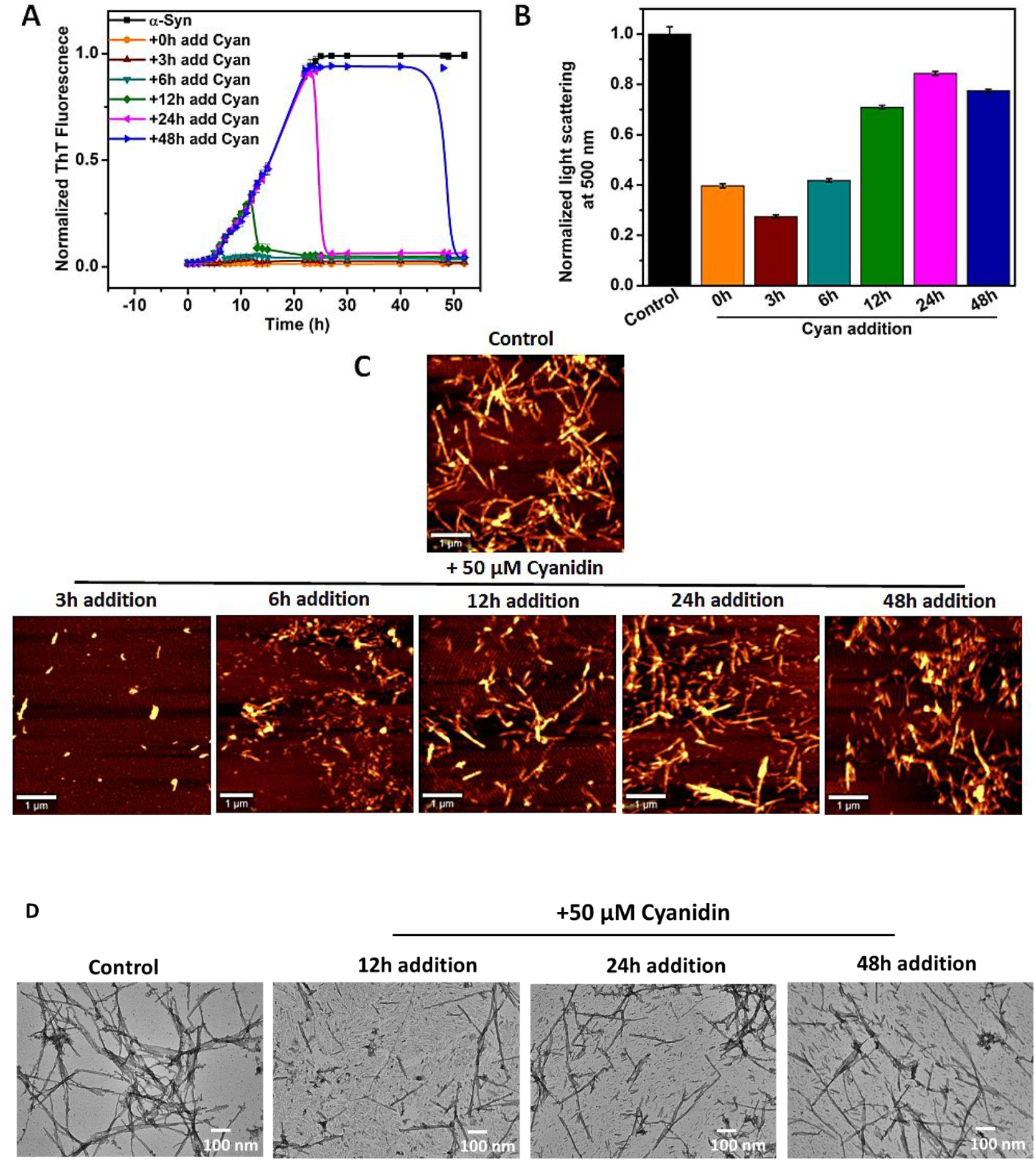
Disaggregation of α-syn amyloid fibrils by Cyanidin (Cyan) (A) ThT fibrillation kinetics of α-syn (70 μM) in the presence of 50 µM Cyanidin (α-syn :Cyan ≈ 1:1), with Cyan added at 0h, 3h, 6h, 12h, 24h and 48h. (B) Normalized Light scattering analysis of α-syn species formed at the end of fibrillation after addition of Cyan at different time-points.

SDS-PAGE results further validated the occurrence of disaggregation of α-syn amyloid fibrils, as disaggregated bands were visualised in the insoluble fraction gel (Figure S3). The insoluble fraction of Cyanidin-treated α-syn samples with Cyanidin added at different stages of the fibrillation pathway (P1, P2, P3, P4 and P5 correspond to the samples where Cyanidin was added to 0, 3, 12, 24 and 48 hours, respectively) was analysed. Disaggregated bands were observed in the P3, P4 and P5 lanes, indicating that Cyanidin was able to act on the fibrils that had already formed during 12, 24 and 48-hour stages of the fibrillation pathway (**Figure S3**).

### Modulation of α-synuclein fibrillation pathway to form SDS-resistant higher order structures in the presence of Cyanidin

To examine the impact of Cyanidin on the population of various α-Syn species formed during aggregation and its influence on the modulation of the α-Syn fibrillation pathway, size-exclusion chromatography (SEC) was utilized wherein higher order structures were separated from the monomeric species corresponding to different peaks. Samples of α-Syn, incubated both with and without Cyanidin (50 μM) during fibrillation, were withdrawn at regular intervals (0, 3, 6, 24, 48, and 72 hours). The soluble fractions obtained after centrifugation were analyzed using SEC at an absorbance of 275 nm. Freshly prepared α-Syn solution showed a major monomeric peak at ∼15.5 ml, corresponding to apparent molecular weights of ∼44 kDa, as calculated from the standard calibration curve (Figure S4). Further, Cyanidin-treated α-Syn samples withdrawn at different intervals of time (Figure 4A) showed a major monomeric peak at ∼15.5 ml and a small oligomeric peak at ∼9.0 ml, corresponding to apparent molecular weights of ∼44 kDa and ∼670 kDa (∼15-mer), respectively. The apparent molecular weight of monomeric α-Syn, instead of its actual molecular weight of 14.4 kDa, is due to the natively unfolded nature of the intrinsically disordered protein. The SEC profiles of α-Syn during aggregation revealed an oligomeric peak when Cyanidin was added at 0 hours and the sample was withdrawn immediately, demonstrating the instant effect of Cyanidin on α-syn aggregation. This indicates that Cyanidin acts immediately on α-syn, altering its conformation from a monomeric to an oligomeric state (Figure 4A). By the end of fibrillation (72 hours), the soluble protein content remained significantly intact, eluting primarily as monomers, with higher-order oligomers present in the void volume. As shown in Figure 4B, Cyanidin was added at different time intervals during the fibrillation pathway (0, 3, 6, 12, 24, and 48 hours) and allowed to aggregate for 72 hours. The results indicate the formation of higher-order structures when Cyanidin was added up to 6 hours into the aggregation pathway, corresponding to peak 1. When Cyanidin was added at later stages, both the monomeric and oligomeric peaks were significantly diminished. This suggests that the addition of Cyanidin to pre-fibrillar or fibrillar species does not induce the formation of oligomers, and the absence of the monomeric peak is due to the conversion of monomers into the fibrillar aggregates.

**Figure 4:**
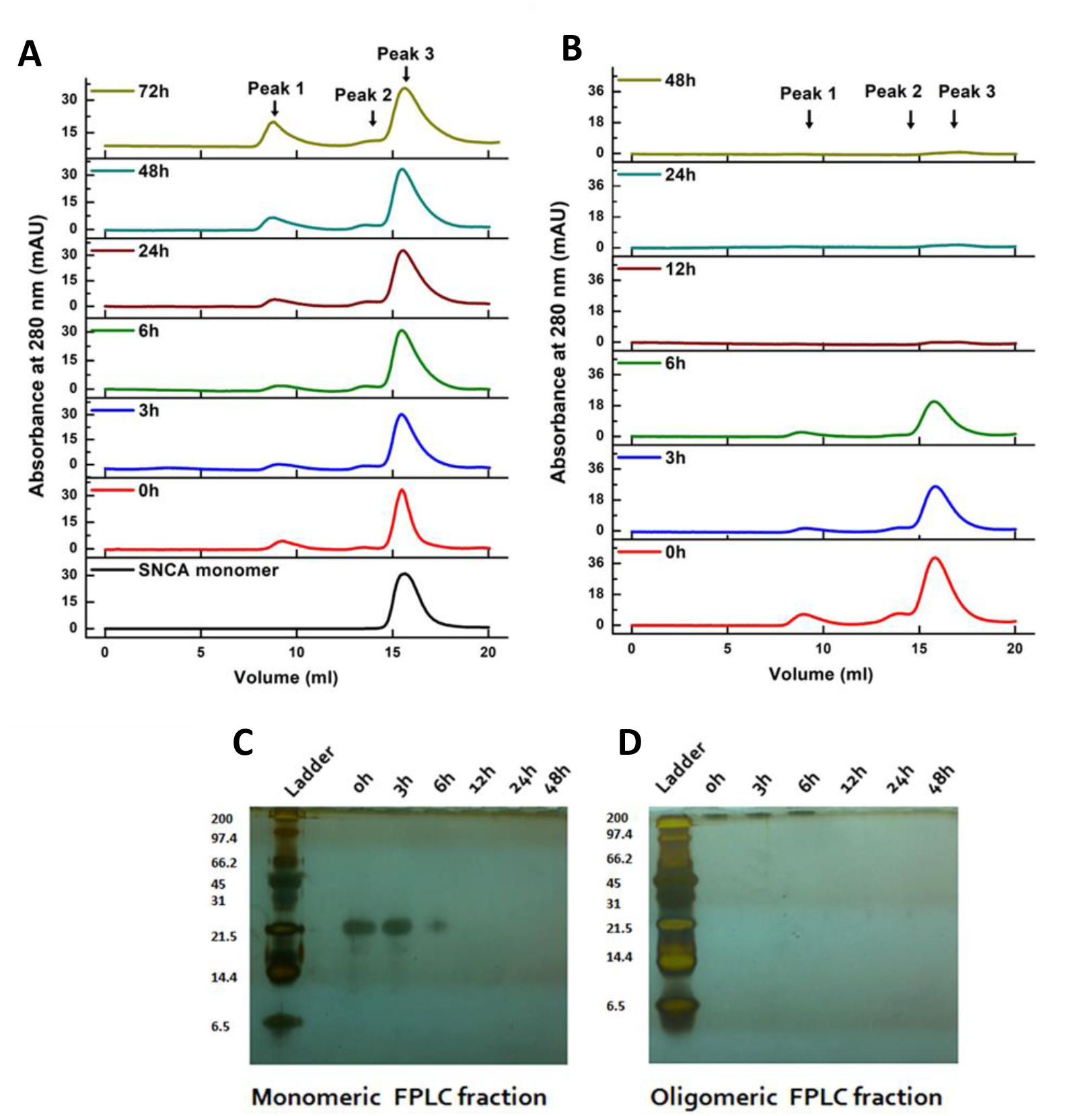
Size-exclusion gel chromatography of α-Syn. Upper panel. **(A)** shows SEC profile of α-Syn samples withdrawn at different time-points (0, 3, 6, 24, 48, and 72h) of fibrillation after addition of 50 µM Cyanidin at 0h of the fibrillation pathway. Monomeric (black) α-Syn elutes at ∼15.5 ml.SEC profile of α-Syn (red), where Cyanidin is added at 0h and the sample is immediately withdrawn at 0h, shows the presence of higher-order species eluting at ∼9.8 ml.along with monomeric species at 15.5 ml. **Upper panel (B)** shows SEC profile of α-Syn samples, where the samples are allowed to fibrillate for 72 hours after addition of Cyanidin at different time-points (0, 3, 6, 12, 24, and 48h) of the fibrillation pathway. The **lower panel** shows two separate SDS-PAGE gels: **(A)** corresponding to Peak 1 (monomeric FPLC fraction of time-point addition samples) and **(B)** corresponding to Peak 3 (higher-order oligomeric FPLC fraction of time-point addition samples).

The soluble eluted samples from FPLC were subjected to two separate SDS-PAGE gels-one corresponding to Peak 1 (monomeric FPLC fraction) (Figure 4C) and the other to Peak 3 (higher-order oligomeric FPLC fraction) (Figure 4D). To determine the nature and stability of the oligomers formed, electrophoresis was performed under denaturing conditions. The gel in Figure 4D revealed the formation of SDS-stable α-syn higher-order structures in the presence of Cyanidin when Cyanidin was added up to 6 hours into the fibrillation pathway, with an apparent molecular weight of ∼200 kDa and above, which were not observed in the untreated α-Syn samples. Cyanidin’s impact on the nucleation phase of α-syn leads to the formation of large droplet-like structures with an average height of ∼20 nm, as observed in the AFM images. This structural transformation likely imparts stability to these structures, making them resistant to denaturation by SDS.

Additionally, the eluted samples from FPLC (corresponding to peak 1) were subjected to dynamic light scattering (DLS) analysis. The DLS data revealed the size of the α-syn aggregates formed in the presence of 50 µM Cyanidin. These aggregates exhibit a hydrodynamic radius of approximately 125 nm. (Figure5B). The hydrodynamic radii of α-syn monomer (Figure 5A, black, 3 nm radius) and fibrils (Figure 5A, green, <100 nm radius) have also been determined by DLS measurements, and match with those reported in the literature. It is important to note that these hydrodynamic radii are rough estimates, assuming a globular shape of the present species.

**Figure 5:**
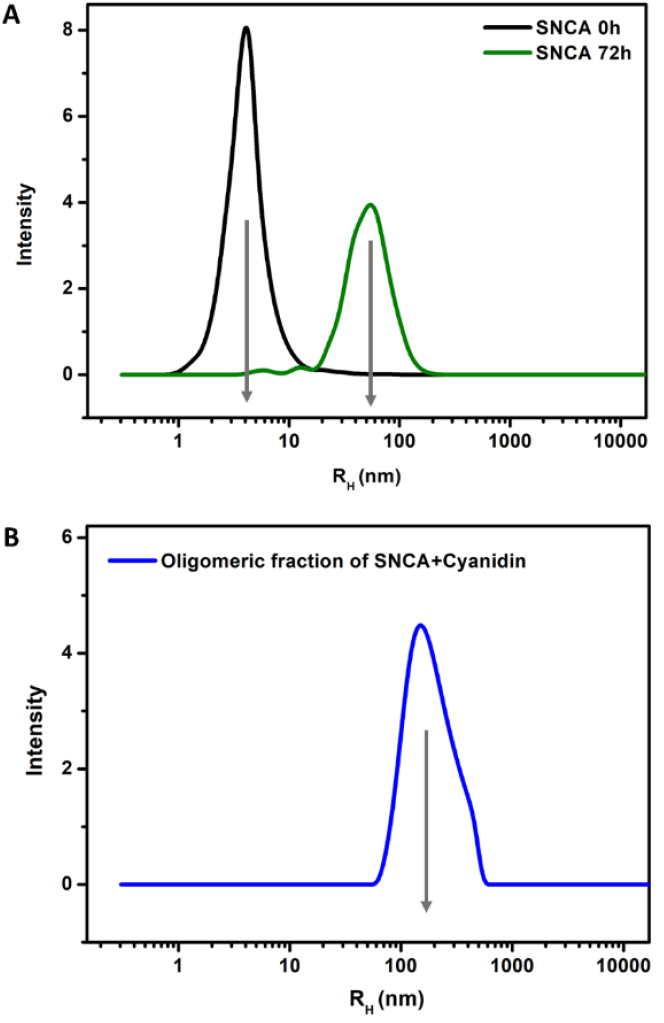
Representative graph of hydrodynamic radius in nm of α-syn species determined using DLS. Graph A shows the hydrodynamic radius (x-axis) of monomer (black) and fibrils (green) and Graph B shows radius of oligomers (blue).

### Structural Characterization

We further investigated the structural transformation of α-syn in the presence of Cyanidin during the aggregation process using CD spectroscopy. Monomeric α-syn exhibits a negative peak at 208 nm in the CD spectrum (Figure 6A), indicating a random coil conformation in solution at 0 hours. Fibrils formed in sodium phosphate buffer, without Cyanidin, display negative ellipticity at 218–220 nm (Figure 6A). Interestingly, the higher order structures formed in the presence of 50 μM Cyanidin exhibit a clear transition from random coil structure to an α-helical form, characterized by two negative ellipticities at 208 and 220 nm. This suggests that, during fibrillation, α-syn undergoes a transformation from random coil conformation to α-helical structure in the presence of Cyanidin.

**Figure 6:**
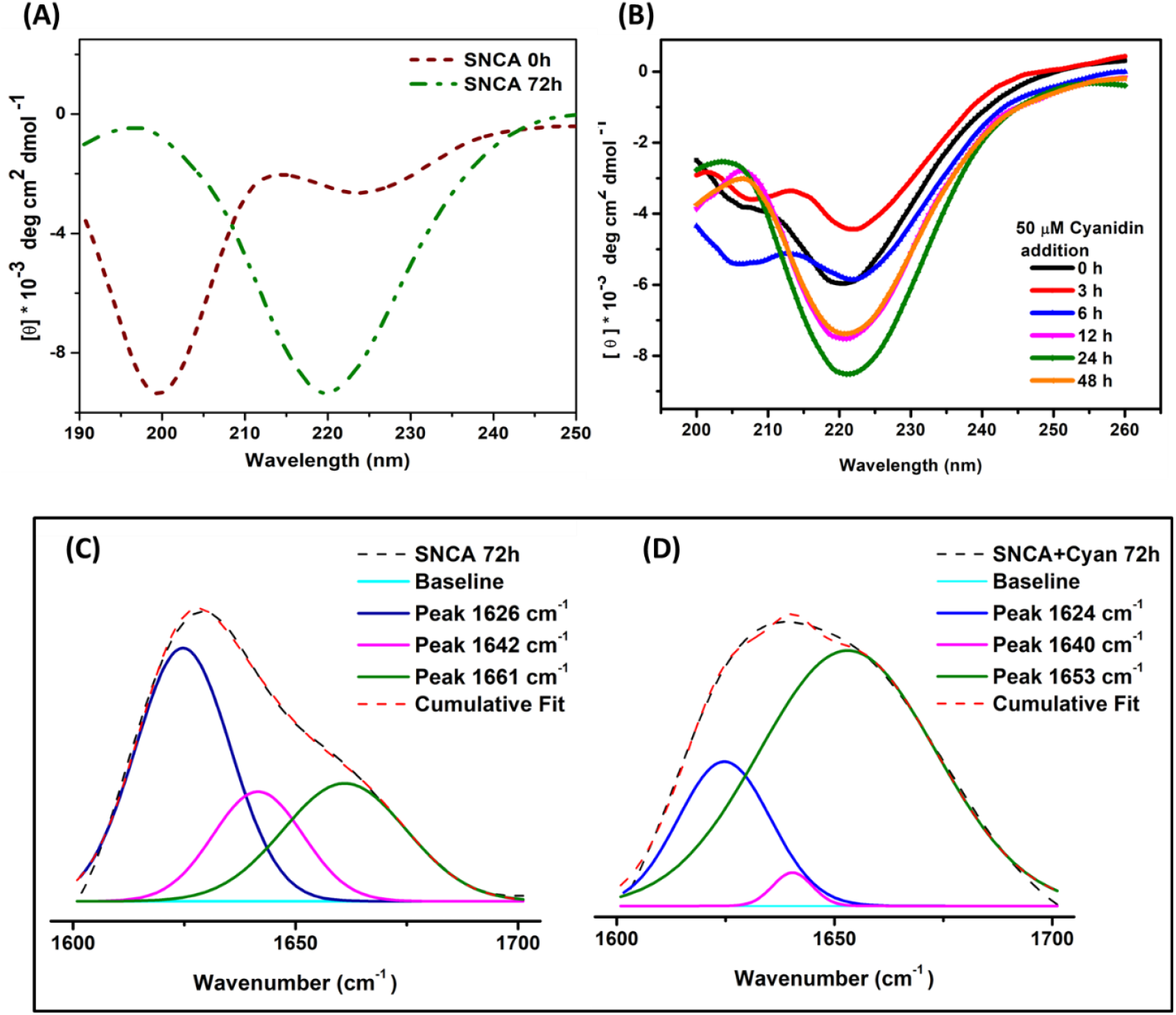
**Amide I region of the FTIR spectra** of control α-synuclein (left panel) and Cyanidin-modified α-synuclein (right panel) taken at the end of the fibrillation process. Dashed black lines are experimental signals; dashed red lines are cumulative fits. Signals under the curve represent deconvoluted experimental curve-fit signals positioned at ∼1624 cm^−1^ (blue lines), ∼1640 cm^−1^ (pink lines), ∼1661 cm^−1^ (green lines).

To confirm the presence of α-helical conformation of the Cyanidin-generated higher-order aggregates, we also performed Fourier transform infrared (FTIR) spectroscopy. By analysing the amide I region (1700–1600 cm^−1^) using FTIR spectroscopy, we extracted secondary structure information of the aggregates. Figure 6C corresponds to α-syn fibrils displaying a spectrum dominated by a band at 1624 cm^−1^ in the second derivative of FTIR spectra. This band is attributed to the presence of beta-sheet structure^15, 16^. Additionally, another band at 1660 cm^−1^ is assigned to various structural elements (α-helix, β turn, random coil or their combination), all of which absorb in this wavelength range. The presence of this band indicates that a portion of the α-syn sequence is not involved in the core β-sheet structure of the fibril ^17^. Native β-sheets in proteins absorb in the 1630–1640 cm^−1^ range, whereas β-structured aggregates typically absorb below 1630 cm^−118^.

When comparing the spectrum of higher order aggregates in the presence of Cyanidin, we observed a decrease in β-sheet content and an increase in α-helical content, as evidenced by the absorption peak at 1653 cm^−1^ (Figure 6D). FTIR investigation strengthens the present finding of distinct helical oligomeric higher-order aggregates in the presence of Cyanidin, characterized by the major absorption peak at 1653 cm^−1^, indicative of the presence of helical conformation.

### Seeding capability

To further explore whether Cyanidin-generated higher order structures serve as templates for α-Syn polymerization, a seeding assay was conducted. We began by aggregating α-Syn to produce amyloid fibrils, which served as a source of seeds upon sonication. The ability of α-Syn amyloid fibrils to seed α-Syn aggregation is reflected in the ThT fibrillation kinetics. Seeding capability is evident as a reduction of the lag phase duration compared to unseeded aggregation, as observed in Figure 7A. The aggregation kinetics of α-syn alone shows a lag phase of 5 hours, followed by an exponential phase, and finally reaching a plateau/stationary phase at 48 hours. Conversely, in the presence of 50 μM Cyanidin, the aggregation kinetics of α-syn were severely affected. The Cyanidin-induced seeds showed a reduced lag time compared to non-seeded α-syn kinetics, but an increased lag time compared to α-syn seeded kinetics. Additionally, when untreated α-Syn seeds were supplemented with Cyanidin (50 μM), α-syn fibrillation kinetics showed a significant reduction in ThT fluorescence. This indicates that Cyanidin effectively inhibits fibril elongation despite the presence of α-syn species with seeding capability. The increased lag time observed with Cyanidin-induced seeds compared with untreated α-Syn seeds indicate that the higher order structures formed in the presence of Cyanidin act as a kinetic barrier for monomer addition and provide hindrance for nucleus formation during fibril polymerization. The Cyanidin-induced seeds are considered ‘on-pathway’ species as they tend to have the capability to polymerize the monomeric α-Syn into fibrils. Previous studies have suggested that the on- and off-pathway identity of oligomers have no direct connection to their cytotoxicity ^19^. Depending on the seeding mechanism, seeds can either be incorporated into the fibril, transferring their morphology to the growing αSyn fibril (seed elongation), or the seed surface can act as a catalyst for the nucleation of new αSyn fibrils (seed surface-mediated nucleation), without necessarily transferring the seed morphology^20^. To investigate the transmission of seed morphology to growing WT αSyn fibrils, it was essential to distinguish between the morphologies of seeded protein fibrils and WT αSyn fibrils using TEM.

**Figure 7:**
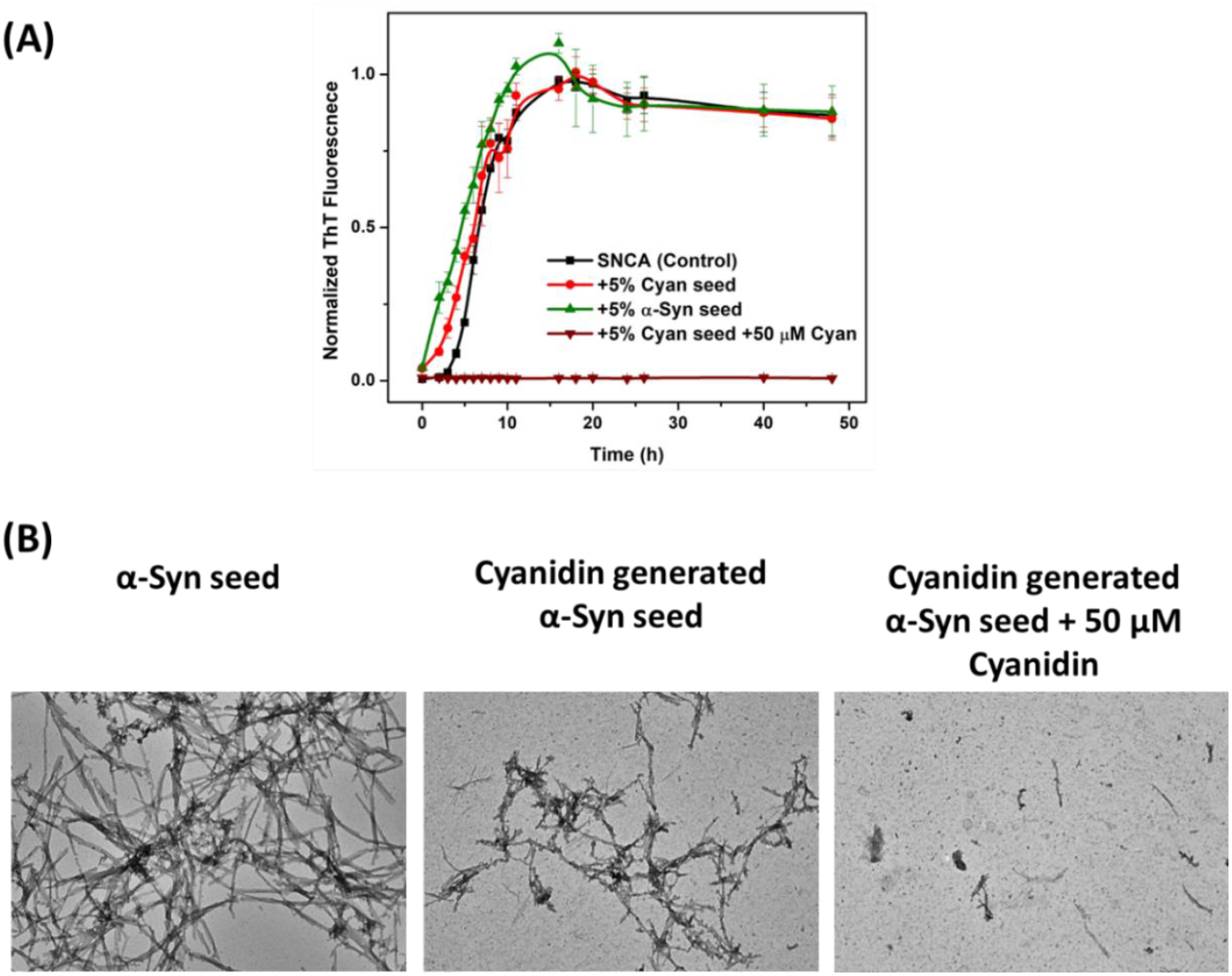
Seeding capability of Cyanidin-induced α-syn seeds. (A) (Upper panel) Fibrillation kinetics of α-Syn in the absence (black) and presence of Cyanidin (50 μM) treated (red) and untreated seeds (green) (5% v/v) shows reduced lag time of fibrillation in the presence of Cyanidin-induced seeds as compared to the untreated α-Syn seeds. Negligible rise in ThT fluorescence in the presence of Cyanidin (50 μM) when added to the Cyanidin-seeded α-Syn fibrillation (green line) shows fibril inhibition. The data have been normalized and the error bars represent ±SD (n = 3). (B) (Lower panel) TEM images showing the morphology of α-Syn species formed in the presence of α-Syn seed, Cyanidin-generated α-Syn seeds and Cyanidin treatment to Cyanidin-generated α-Syn seeds (left to right). Scale bars, 100 nm.

Using negative stain transmission electron microscopy, we further confirmed the fibrillation of α-syn monomer with Cyanidin generated α-syn seeds. In the presence of Cyanidin, these seeds prevented the formation of α-syn fibrils, in contrast to the absence of Cyanidin, where only a small number of fibrils were accumulated (Figure7B). Cyanidin was seen to slightly modulate α-syn fibrillation in the presence of Cyanidin induced seeds, whereby the treated products were less abundant, and appeared more diffused as compared to the tightly-packed appearance of the untreated fibrils. Thus, seeding studies revealed that the Cyanidin-induced higher order structures are ‘partial on-pathway’ species that fail to build into mature elongated α-syn amyloid fibrils in the fibrillation pathway. Although the presence of Cyanidin-induced seeds significantly reduces the lag phase of αSyn fibril formation, the morphological features of the resulting αSyn amyloid fibrils remain unchanged, suggesting a seed surface-mediated nucleation mechanism.

ANS (1-anilino-8-naphthalenesulfonic acid), a commonly employed solvent-sensitive fluorescent dye, detects intermediates containing solvent-exposed hydrophobic clusters during the formation of β-sheet fibrils along the fibrillation pathway. ANS can characterize proteins by revealing their hydrophobic nature. When ANS is bound to proteins, it exhibits blue-shifted fluorescence maxima and increased fluorescence intensity, changes attributed to protein’s hydrophobicity. Figure 8 (right panel) shows ANS fluorescence of α-syn alone and with Cyanidin at the monomeric and fibrillar stages. Maximum ANS fluorescence was observed for α-syn fibrils, while minimal or reduced fluorescence was seen for monomeric α-syn control. These observations suggest the presence of exposed hydrophobic patches on α-syn fibrils compared to the monomeric protein.

**Figure 8:**
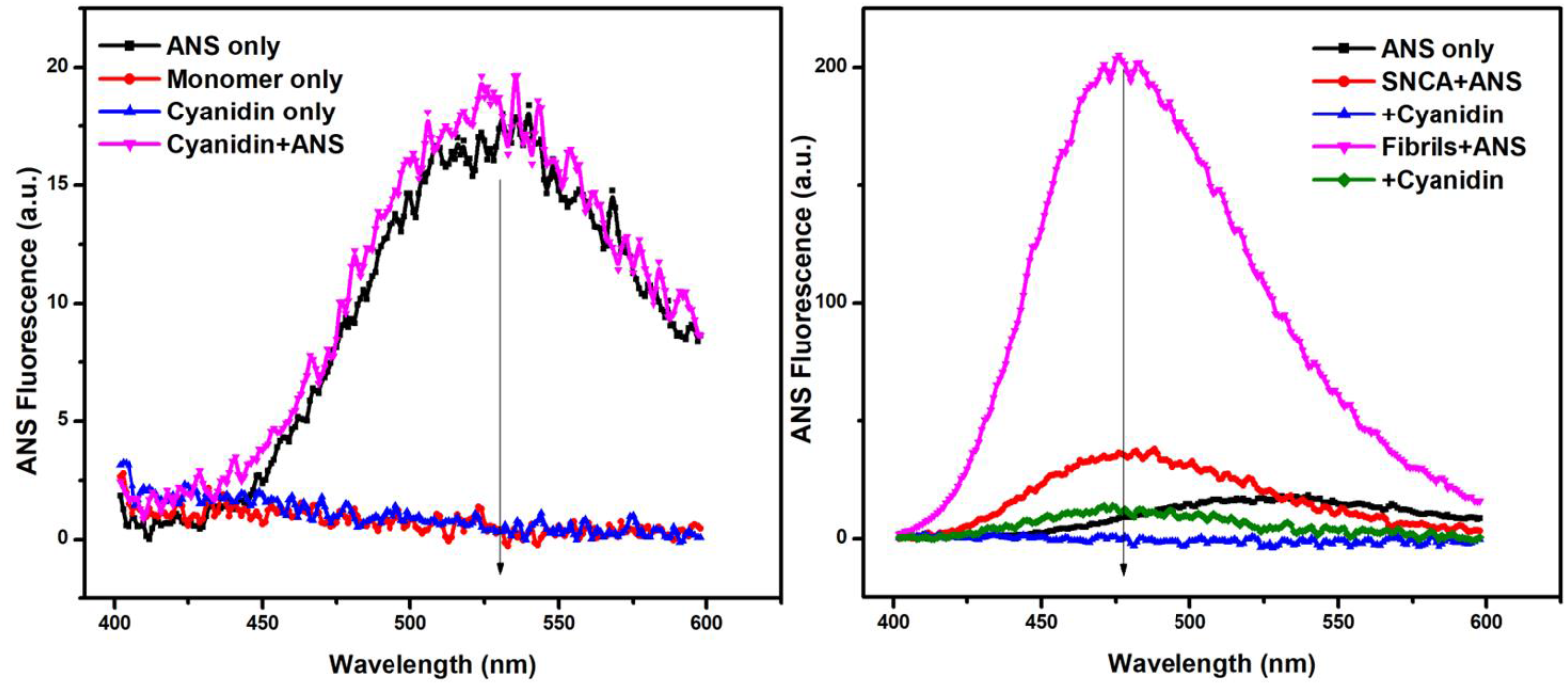
Hydrophobicity of α-syn species in the presence of Cyanidin (50 µM). (Left panel) Emission spectra of ANS alone, ANS binding to monomeric α-Syn, Cyanidin alone and ANS binding to Cyanidin. (Right panel) ANS binding to monomeric α-syn in the presence of Cyanidin and ANS binding to aggregating species of α-Syn in the absence and presence of Cyanidin.

Further, ANS fluorescence was analysed by adding 50 μM Cyanidin to the monomeric and fibrillar stage (i.e., 0h and 48h) of the α-syn fibrillation pathway. Negligible ANS fluorescence was observed for both samples where Cyanidin was added at the initial and stationary stages of the fibrillation process. When cyanidin was added at the initial stage of the fibrillation, the observations suggest that most of the exposed hydrophobic patches were likely utilised for higher order structure formation and thus were unavailable for binding with ANS dye. However, low fluorescence in the case where Cyanidin was added at the stationary stage suggests that the hydrophobic patches could be buried or masked by Cyanidin.

To assess whether Cyanidin’s inhibitory effect on α-syn aggregation is sufficient to protect cells from α-syn-mediated cytotoxicity, we examined the cytotoxicity of α-syn aggregates produced in the absence or presence of Cyanidin using MTT-based cell viability test (Figure 9). We observed that incubating SHSY-5Y cells with α-syn fibrils significantly reduced cellular viability. However, adding Cyanidin-treated α-syn samples resulted in approximately a 10% increase in cell viability compared to those exposed to the α-Syn fibrillar solution. This suggests that Cyanidin-mediated inhibition of α-syn fibrillation reduced the toxicity of the α-syn sample. Further, we also analysed the toxicity of other samples where Cyanidin was added at different time intervals during the fibrillation pathway. Notably, by measuring the metabolic activity with the MTT assay, we demonstrated that treating SHSY-5Y cells with different species of α-syn, generated after 72hours of fibrillation, resulted in variable toxicities. The α-syn species generated after the addition of Cyanidin at the initial stages of the fibrillation pathway (0, 3, and 6 hours) were observed to be less cytotoxic compared to the α-syn fibrils. In contrast, the addition of Cyanidin at the later stages (24 and 48 hours) produced toxic species similar to that of control α-syn fibrils.

**Figure 9:**
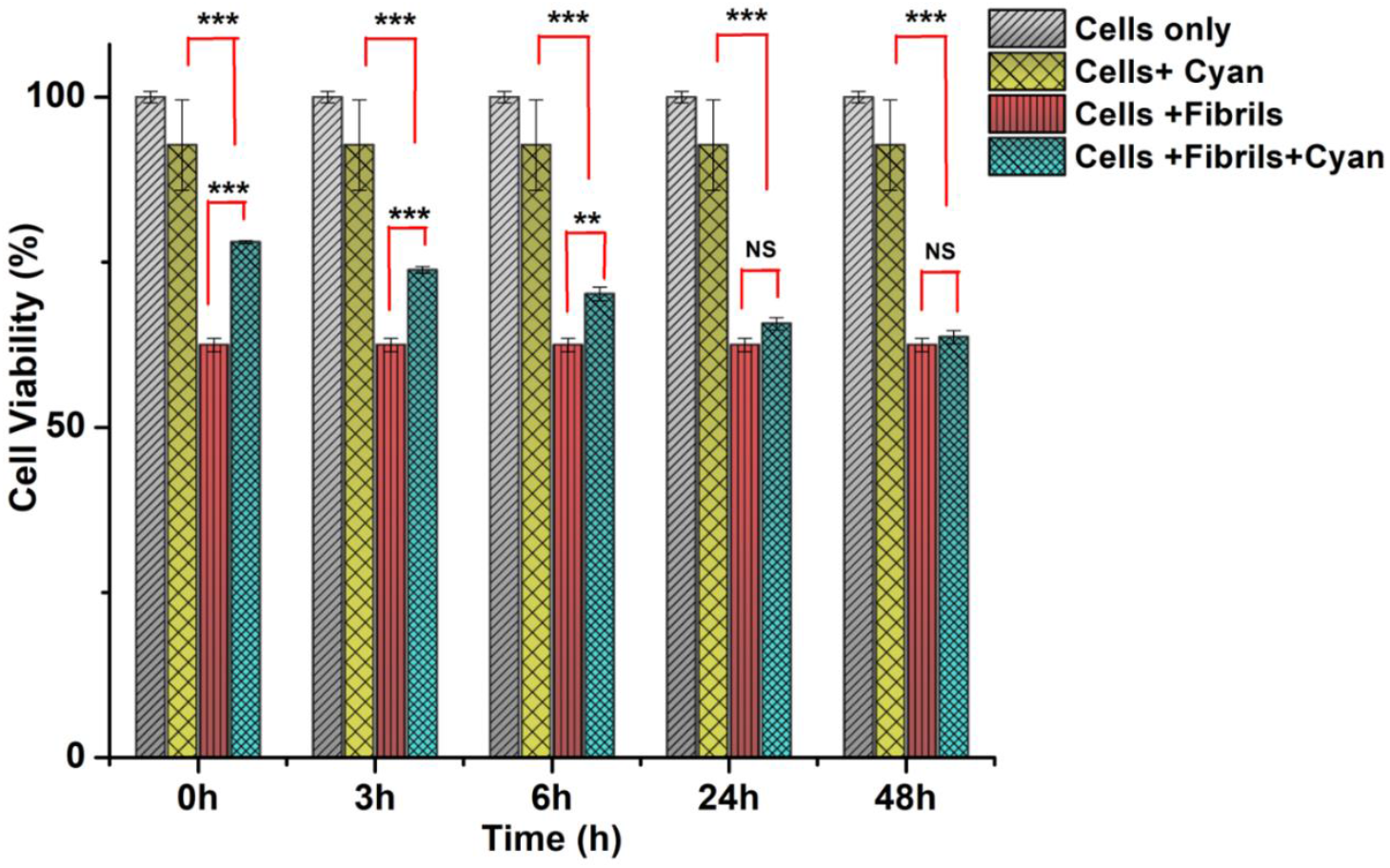
MTT cytotoxicity assay for analysing the cytotoxic effect of Cyanidin treated and untreated α-syn aggregates on the viability of SH-SY5Y cells. MTT assay reveals an increased viability of SHSY5Y cells in the presence of Cyanidin-induced aggregates, when Cyanidin was added at 0, 3, and 6hours of α-syn fibrillation pathway and no significant change in the toxicity of the aggregates was observed when Cyanidin was added at 24 and 48 hours of the α-syn fibrillation pathway. Values represent ± SD.The statistical analysis was done using Student’s t test.Triple asterisk *** indicates P < 0.005

### Cyanidin binds weakly to α-Syn

To obtain reliable values for Cyanidin−α-syn dissociation constants and explore the accessibility of Cyanidin to the tyrosine’s present in α-Syn, we monitored the tyrosine fluorescence of α-Syn in the presence of increasing concentration of Cyanidin (2–30 μM). Excitation was performed at 275 nm, and emission intensity was recorded from 280 to 400 nm. A concentration-dependent decrease in tyrosine fluorescence indicated an interaction between α-Syn and Cyanidin (Figure 10A). Using the Stern–Volmer quenching equation, the bimolecular quenching constant (*k*_q_) was determined to be more than 10^10^ M^−1^ s^−1^ (*k*_q_ = 2.0 × 10^13^ M^−1^ s^−1^) suggesting a static quenching mechanism and the formation of a non-luminescent ground-state complex between Cyanidin and α-Syn. The binding parameters were assessed using the modified Stern– Volmer equation, which showed a linear relationship between fluorescence quenching intensity and Cyanidin concentration, as depicted in the plot of log (Fo–F/F) versus log (Q) (Figure 10B). The results indicated that α-Syn binds to Cyanidin at an approximately equimolar ratio of 1:1. A binding constant (K_a_) of 8.4×10^5^ M^−1^was determined at 37°C, suggesting that weak non-covalent interactions play a role in the Cyanidin-mediated inhibition of α-Syn fibrillation. We also used ITC to quantify the interaction between Cyanidin and monomeric α-syn by measuring the rate of heat produced from the binding as the protein was titrated with the ligand (Figure 10C, upper panel). ITC binding experiment between Cyanidin and α-syn were performed at 37^°^C, where α-syn was used as titrant in the cell at a concentration of approximately 70 μM, and Cyanidin at approximately 10-fold higher concentration as titrant in the syringe. The resulting binding isotherm reflecting the integrated heat of reaction (Figure 10C, lower panel) was well fit by a model consisting of one binding site (stoichiometry of 1.0). The dissociation constant (K_d_) derived from the binding isotherm was relatively moderate, measured in mM range (3.76 x 10^-5^). The dissociation constant (K_d_) derived from the modified Stern–Volmer plot from steady-state fluorescence studies also indicated a weak binding affinity between Cyanidin and α-Syn, showing the significance of both techniques in analysing the binding parameters. The results indicate that weak non-covalent interactions play a significant role in the binding of Cyanidin to α-Syn. IDPs are known to interact with their targets with high specificity and low binding affinity, enabling precise ligand binding and playing a crucial role in signal transduction ^21^. Therefore, weak binding of Cyanidin to α-syn and significant suppression of fibrillation, highlights the importance of weak interactions in the modulation of α-syn fibrillation.

**Figure 10:**
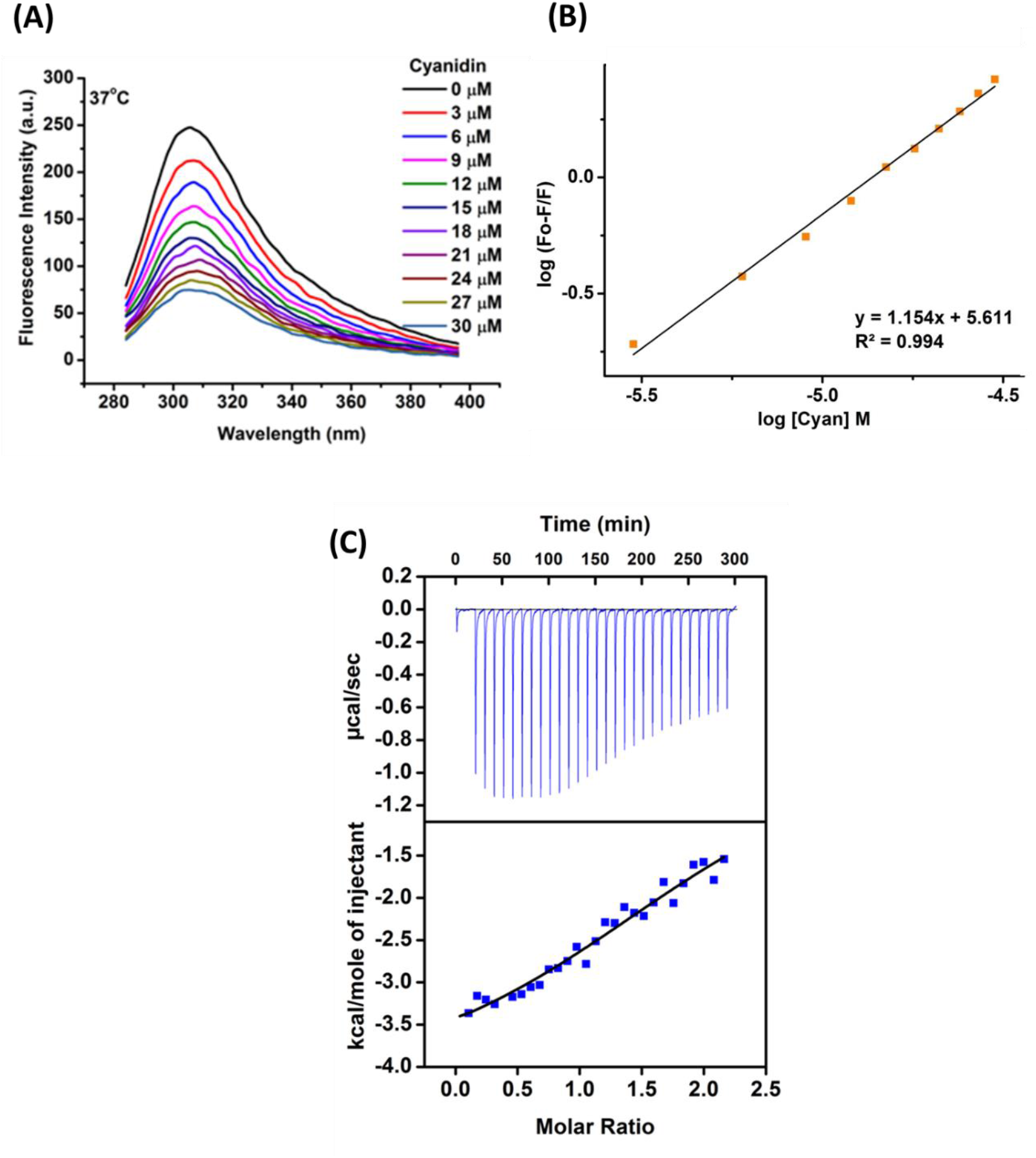
Steady-state fluorescence and ITC study show moderate mode of binding between Cyanidin and α-Syn. (A) Steady-state fluorescence shows continuous quenching of intrinsic tyrosine fluorescence of α-Syn (0.3 mg/mL) upon titration with an increasing concentration of Cyanidin (2–30 μM) at 37°C. (B) Modified Stern–Volmer plot of log (F_0_– F)/F versus log [Q] shows moderate binding interaction between α-Syn and Cyanidin (K_a_ = 8.4×10^5^ M^−1^). (C) An isothermal titration calorimetry study shows moderate interaction between Cyanidin and α-Syn at 37°C using a α-Syn to Cyanidin ratio of 1:10. The upper panel of the results displays the raw data plot of heat flow over time during the titration, while the lower panel shows the total normalized heat released as a function of ligand concentration. The solid line represents the one-site fit for the obtained data

After examining the modulation of molecular mechanism of α-Syn fibrillation by Cyanidin and determining its binding affinity with monomeric α-Syn, we aimed to identify the potential binding sites of Cyanidin on α-Syn. Molecular docking was conducted on the α-Syn structural conformations (RMSD cutoff 19Å) derived from structural ensembles obtained through NMR and PRE measurements ^22^ from the Protein Ensemble database (pE-DB) ^23^, as shown in our previous work. The docking studies indicated a significant binding affinity of Cyanidin to all the selected α-syn conformations, with a binding free energy of ∼7.0 kcal/mol (**Figure 11 and Table S1**). Cyanidin primarily binds to Lysine residues through hydrogen bonds, in regions spanning the N-terminal and NAC region,is shown in **Figure 11**. Additional other non-polar interactions also contribute to Cyanidin binding to the protein. Hydrogen bonding of Cyanidin to α-syn is specifically observed in the NAC region, particularly between residues 64-110. Given that the NAC region is crucial for α-syn aggregation ^24^, the preferential binding of Cyanidin to this region is likely to prevent the aggregation-prone NAC region from participating in the usual fibrillation process, resulting in the formation of less hydrophobic higher-order species. Although molecular docking, a structure based drug discovery approach ^25^, has limitations for studying interactions of intrinsically disordered proteins, it can still provide insights into the potential binding sites of Cyanidin on α-syn.

**Figure 11:**
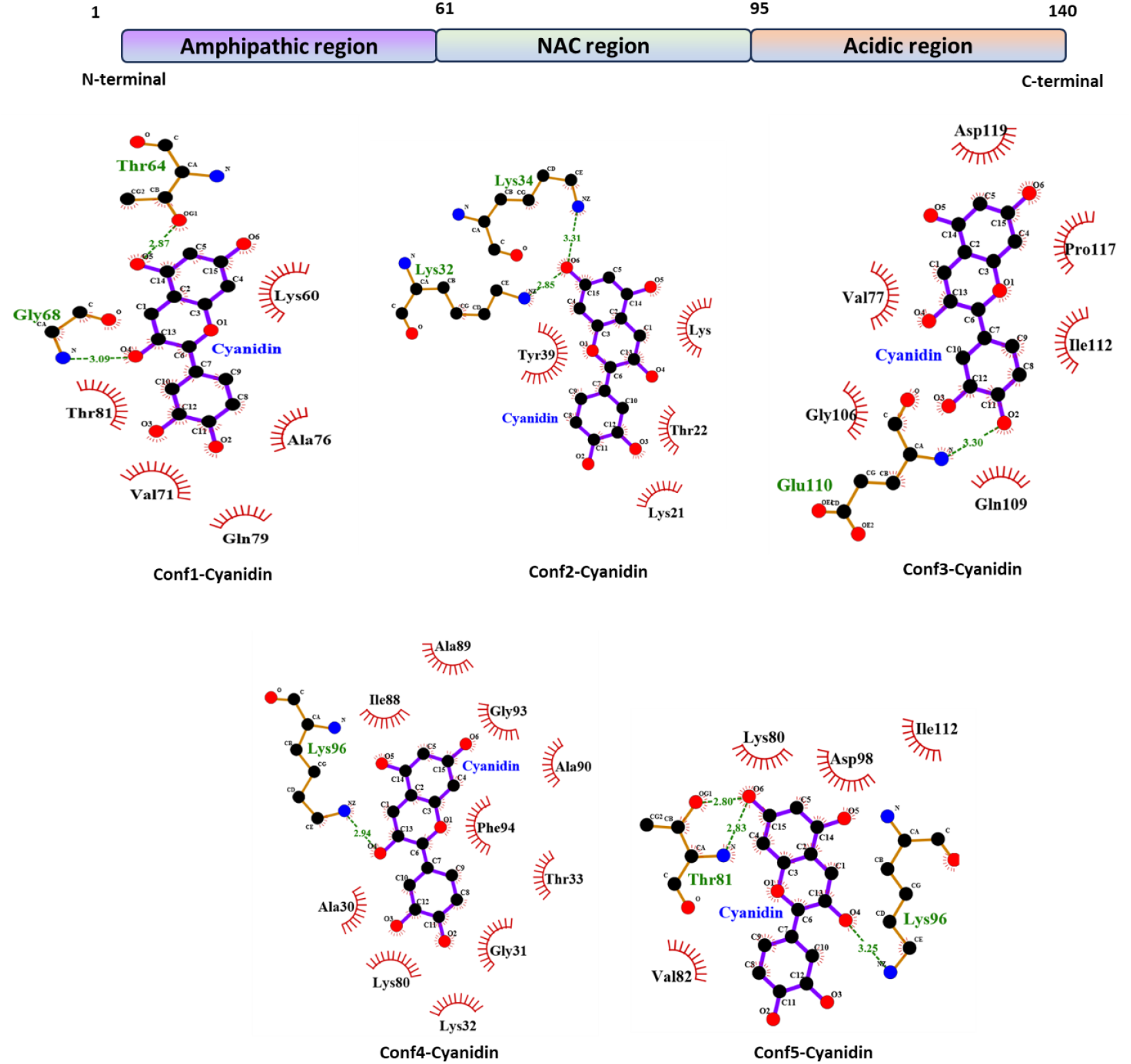
Molecular docking of Cyanidin on α-synuclein. Schematic of α-synuclein (α-syn) and molecular docking analysis of five α-syn conformations (conf1-5) with Cyanidin. The study reveals preferential binding sites in the N-terminus and NAC region. The interactions were visualized using Discovery Studio.

## Discussion

Understanding the Parkinson’s disease initiation and progression remains challenging due to gaps in our knowledge. This lack of understanding has hindered drug discovery efforts. One promising avenue is inhibiting the transition from liquid to toxic fibrillar phases. Our work investigates how the natural neuroprotector Cyanidin modulates α-Syn fibrillation towards the formation of non-toxic aggregates. To assess the inhibition of α-Syn fibril formation by Cyanidin, ThT fluorescence assay and atomic force microscopy (AFM) experiments were conducted. A 70 μM solution of α-Syn, both with and without Cyanidin treatment, was dried on a mica surface, and AFM images were captured. In the absence of Cyanidin, long fibrillar structures were evident (Figure 3C, control). However, with the addition of 50 μM Cyanidin, large droplet like higher-order structures were visible. Further, upon visualising the Cyanidin treated α-syn samples by TEM, we observed the formation of hollow droplet shaped solid mesh/gel-like structures, containing small protofibrillar species at the periphery of the surface.

It has been shown that α-syn can irreversibly capture cellular components through liquid-liquid phase separation, driving its self-assembly into solid aggregates and ultimately leading to various pathological processes ^26, 27^. Our results specifically demonstrate that Cyanidin induces the formation of liquid-like droplets, which eventually undergo a liquid-to-solid transition to form an amyloid hydrogel containing small protofibrillar species. Cyanidin-induced spherically-shaped higher order α-syn aggregates exhibited significant molecular rigidity (Figure 2). This observation aligns with a study showing a liquid-to-solid-like transition of droplets with liquid-like properties in the presence of molecular crowding agents such as PEG ^28^. Furthermore, we show that these Cyanidin-induced liquid-like droplets are SDS-stable and non-toxic but acquire seeding capability when dysregulated by external forces like sonication. According to the structural analysis, both the N-terminus and NAC regions are identified as the major contributors to phase separation ^29^. Our docking study reveals that Cyanidin binds to both the N-terminus and NAC region, which may explain the stability of liquid-like droplets into hydrogel. A similar scenario has been observed in a study where α-syn, upon interacting with the natural molecule LL-III, stabilizes the droplets and prevents their conversion into a fibrillar state ^30^.

The secondary structure and surface-exposed hydrophobicity are interrelated factors that influence aggregate toxicity. Aggregates with higher β-sheet content and larger hydrophobic areas tend to be the most toxic. To investigate this, ANS was employed to assess the hydrophobicity of the species formed which allowed the comparison of the negligible hydrophobicity of Cyanidin-treated aggregates with that of control fibrils. The immediate interaction of Cyanidin with monomeric α-syn facilitated structural reorganization into species with hydrophobic areas facing each other in an enclosed, hollow-shaped structure (Figure 2). This resulted in the formation of large, droplet-shaped structures with no exposed hydrophobic patches to bind to ANS, providing insights into the surface properties of these droplet-shaped aggregates.

To investigate the interaction between Cyanidin and monomeric α-Syn, we measured the binding affinity using steady-state fluorescence and ITC (isothermal titration calorimetry) studies. The data revealed a weak interaction between Cyanidin and the monomeric form of α-Syn, effectively preventing its transformation into the fibrillar state.

α-synuclein (α-syn) undergoes a critical conformational transition from a disordered state to a β-sheet-rich structure, which plays a key role in the formation of amyloid fibrils. The accumulation of β-sheets and amyloid nature of the assemblies formed in the absence of Cyanidin were analyzed using CD spectroscopy (Fig. 5A). The transition from a disordered structure to β-sheet-rich structures, which occurred without Cyanidin, was indicated by the appearance of a peak minimum around 218 nm ^31^. Interestingly, incubation with Cyanidin inhibited these conformational changes, resulting instead in α-helical rich structures. These structural alterations likely arose from specific interactions between Cyanidin and distinct regions of α-syn, disrupting the forces driving the transition toward a β-sheet-rich state. To validate the CD data on the secondary structure of the Cyanidin induced higher-order species and to confirm the amyloid nature of the control fibrils, FTIR spectroscopy was also employed (Fig. 5C). The amide I band (1700–1600 cm^−1^) is primarily attributed to the C = O stretching vibration due to hydrogen bonding and distortion of amide linkages^32^. Our FTIR results were consistent with the CD findings, showing an accumulation of α-helical rich structures in the presence of Cyanidin.

Computational analysis showed that myricetin interacts with the surface of the β-sheet via H-bonding, weakening the interstrand hydrogen bonds and eventually disrupting the outer layer of the aggregate ^33^. Cyanidin appears to act in an unprecedented way by trapping α-syn in a thermodynamic sink in the form of non-toxic Cyanidin stabilized oligomers. This inhibition property of Cyanidin mainly stems from the weak to moderate preferential interactions of Cyanidin with residues located at the N-terminus and NAC region of α-syn. Cyanidin efficiently inhibits both aggregation and cytotoxicity of α-syn at sub-stoichiometric concentrations and, in addition, is able to convert monomeric α-syn into thermodynamically stable nontoxic Cyanidin-stabilized higher-order structures. Notably, our results also suggested the importance of the anthocyanidin moiety, rather than the sugar moiety.

## Conclusions

Interfering with α-Syn fibrillation has been envisioned as one of the promising disease-modifying approaches for the treatment of PD. Here, we have described Cyanidin as a small molecule natural inhibitor and modulator of α-Syn aggregation towards non-toxic aggregates. We have performed a detailed in vitro biophysical characterization of the modulatory and inhibitory activities of Cyanidin and tested its toxicity in human neuronal cells. Our study demonstrates that Cyanidin interacts effectively with liquid phase droplets of α-Syn. Acting as weak cross-linkers between α-Syn molecules, Cyanidin stabilizes the droplet condensates and prevents their transformation into the fibrillar amyloid state. This stabilization is achieved by increasing the droplet stability and raising the activation free energy barrier required for the transition to the fibrillar state. The non-toxicity of Cyanidin was confirmed in a human cell line. We believe that investigating natural neuroprotective compounds like Cyanidin that interact with specific liquid condensates, such as those of α-Syn, will enhance our understanding of complex disease mechanisms arising from these condensates and may open up new avenues for therapeutic intervention. This work provides detailed insights into liquid to solid transition of natively unstructured α-Syn and its conversion to a stable, non-toxic hydrogel matrix comprising of protofibrillar structures – a non-disease-associated aggregated state, which is highly relevant in Parkinson’s disease pathogenesis.

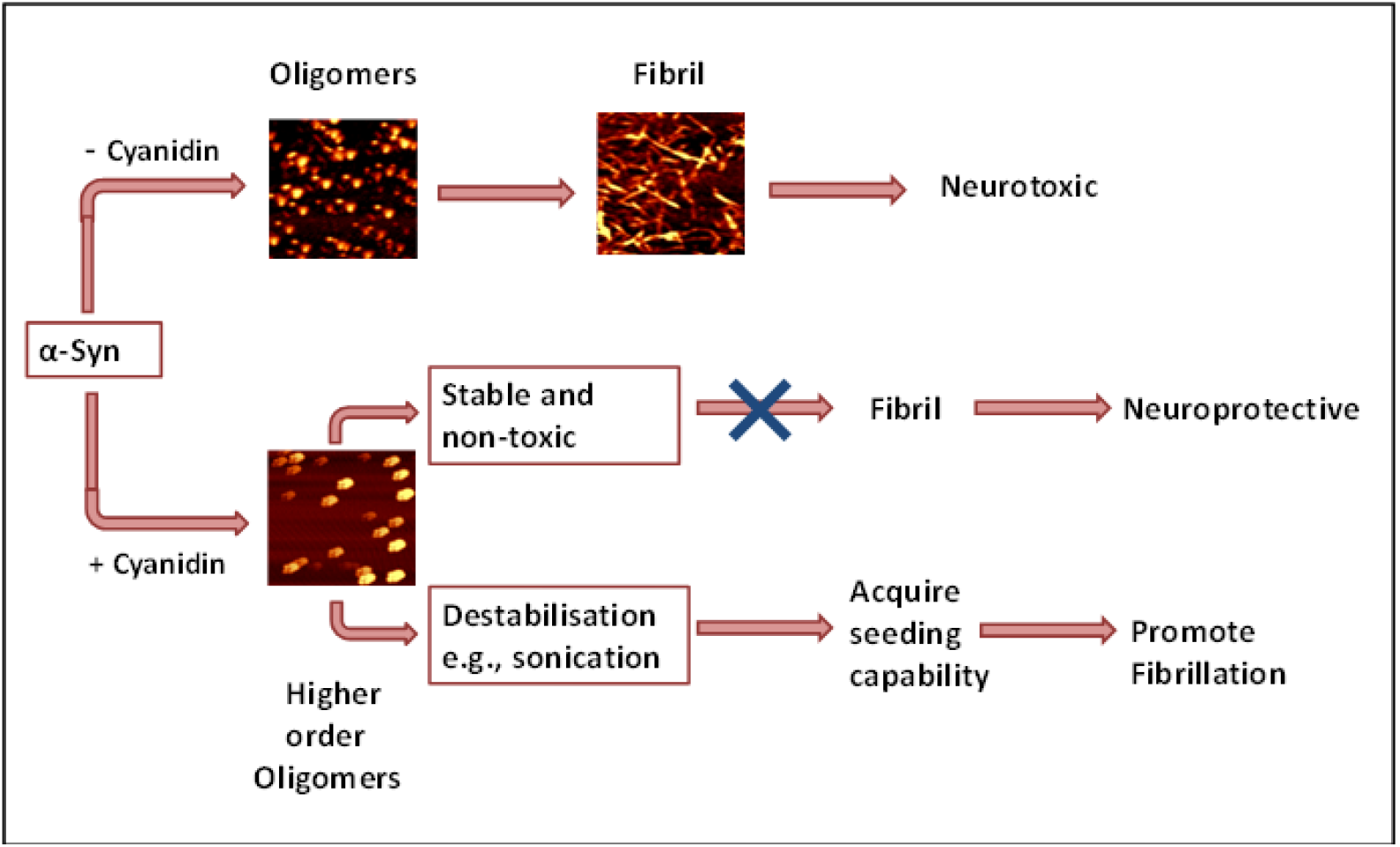
Schematic representation of Cyanidin-mediated suppression of alpha synuclein aggregation.

## Materials and Methods

### Materials

Cyanidin was purchased from Extrasynthese (France). ThT and MTT were from Sigma and rest other chemicals were from Sisco Research laboratory Pvt. Ltd. (SRL) of analytical grade quality. A full length α-syn plasmid was a gift from Dr. Peter T. Lansbury (Harvard Medical School, Cambridge, MA).

### Protein expression and purification

The expression and purification of α-Syn was carried out as described previously^34^ with slight modifications ^35, 36^. The purified protein was resuspended in 10 mM sodium phosphate buffer, pH 7.4 and the concentration was measured at 280nm, with extinction coefficient ε = 5960 cm^-1^. The protein purity was determined by Mass Spectrometry.

### ThT fluorescence assay

The α-Syn fibrillation process was monitored using Thioflavin T (ThT). ThT stock was prepared in 20 mM sodium phosphate buffer at pH 7.4, and Cyanidin was prepared in 100% DMSO. The concentrations of ThT and Cyanidin were measured using molar extinction coefficients of ε = 35,000 M^−1^ cm^−1^ at 412 nm and ε = 4,778 M^−1^cm^−1^ at 544 nm, respectively. α-Syn (70 μM) was mixed with increasing concentrations of Cyanidin (5 to 200 μM) in the fibrillation buffer containing 20 mM sodium phosphate buffer, 100 mM NaCl, at pH 7.4, and allowed to fibrillate for 72 hours under shaking conditions at 200 rpm in a 96-well Corning flat-bottom plate at 37°C. Each well of the 96-well plate contained 200 μl of the sample along with a 3 mm glass bead. A control sample without Cyanidin was also prepared. The final concentration of DMSO in all samples was 1%. Changes in ThT fluorescence were measured using a Varioskan Flash microplate reader (Thermo Scientific) at 480 nm, with excitation at 445 nm and both excitation and emission slit widths set to 5 nm. Measurements were performed in triplicates. The apparent fibrillation rate and lag time were determined by fitting the fibrillation kinetics of α-Syn to a sigmoidal curve using a previously reported empirical formula^37^. For time-dependent addition of Cyanidin and performing disaggregation assay, 50 μM Cyanidin was added at 0, 3, 6, 12, 24, and 48 hours of α-syn fibrillation pathway, and ThT fluorescence was monitored.

### Seeding experiment

The seeds were obtained by incubating α-syn (70 μM) with and without 50 μM Cyanidin for 72 hours at 37°C, under shaking conditions at 200 rpm. The aggregated samples were centrifuged at 14,000 rpm for 30 minutes, and the treated and untreated aggregates were washed twice with phosphate buffer, followed by homogenization by vortexing in 500 μl of 20 mM sodium phosphate buffer, 100 mM NaCl, pH 7.4. The α-syn seeds (5% v/v) formed in the absence and presence of 50 μM Cyanidin (5% v/v) were added to monomeric α-syn (70 μM) along with 20 μM ThT and allowed to fibrillate for 72 hours under shaking conditions. The change in ThT fluorescence was measured using a Varioskan Flash microplate reader at 480 nm with slit widths of 5 nm. The final concentration of DMSO in all samples was 1%.

### ANS fluorescence assay

The samples for The ANS (8-Anilinonaphthalene-1-sulfonate) fluorescence measurements were prepared by adding 50 μM Cyanidin to monomeric α-syn (70 μM) at different time-points (0, 3, 6, 12, 24, 48 hours) of the fibrillation pathway and allowed to fibrillate under the conditions mentioned in ThT section. Before adding ANS, the samples were diluted in the fibrillation buffer to a final protein concentration of 7 μM and mixed with 35 μM ANS (Protein:ANS ratio = 1:5). The ANS spectra were recorded on Cary Eclipse spectrofluorimeter (Varian Inc., California, USA) by exciting at 350 nm. The ANS stock was prepared in water, and its concentration was measured at 350 nm with molar extinction coefficient of 4950 M^−1^ cm^−1^.

### Rayleigh scattering

Samples for Rayleigh scattering measurements were prepared by incubating different concentrations of Cyanidin (5-50 μM) with 70 μM α-syn at 37 ^o^C, 200 rpm for 72 h. DMSO concentration (1%) was constant in all the samples. Static light scattering was recorded at 500 nm, using 1 cm path length cuvette in Cary Eclipse spectrofluorimeter (Varian Inc., California, USA), with excitation and emission slit widths of 2.5 nm.

### Transmission electron microscopy

Electron microscopy was employed to visualize the morphological changes in α-syn aggregates formed due to the addition of 50 μM Cyanidin to the intermediate species of α-syn during its fibrillation pathway. Time-point samples were placed onto carbon-coated 200 mesh copper grids, negatively stained with 1% uranyl acetate, and excess solution was blotted with filter paper. The samples were then dried for 30 minutes and analyzed using a JEOL TEM 2100 microscope operating at an accelerating voltage of 200 kV.

### Atomic force microscopy

To examine the morphology of aggregates and fibrils formed in the absence and presence of 50 μM Cyanidin, samples were diluted in a 20 mM sodium phosphate buffer containing 100 mM NaCl at pH 7.4, using a ratio of approximately 1:3. A 20 μl aliquot of the diluted samples was then loaded onto freshly cleaved mica blocks. A freshly cleaved mica sheet was affixed to a microscope slide, and the mica surface was exfoliated twice before applying the sample. These mica blocks were washed with deionized water and allowed to dry before imaging. Images, consisting of 512 x 512 data points, were captured using an AFM (WITec GmbH, Germany) operating in non-contact mode. The cantilevers used had a force constant of 40 N/m and a resonance frequency of 75 kHz, with a constant scan rate of 0.5 Hz. Image analysis was performed using Project FOUR software.

### SDS-PAGE

α-Syn samples treated with 50 μM Cyanidin at different time points 0 h, 24 h, and 48 h as well as the disaggregation samples were prepared similarly as for size-exclusion studies. The samples were ultracentrifuged at 100000g (Sorvall MTX 150 ultracentrifuge, Thermo Scientific, S52-ST rotor) for 30 minutes. The fractions were separated and resolved on two separate gels - soluble and insoluble – in 0.75 mm-thick plates at 80 V. A broad range protein standard (6.5–210 kD) (Bio-Rad, CA, USA) was used as the molecular weight marker.

### Circular dichroism spectroscopy

To determine the secondary structure, 50 μM Cyanidin was added to the intermediate species in the α-syn fibrillation pathway. The samples were prepared in a 20 mM sodium phosphate buffer with 100 mM NaCl at pH 7.4, incubated at 37°C for 72 hours at 200 rpm, maintaining a constant DMSO concentration across all samples. Far-UV CD spectra were recorded between 190 and 260 nm at 25°C using a J-815 spectropolarimeter (Jasco, Japan) equipped with a Peltier device. The spectra were obtained using a 0.1 mm quartz cuvette at a scanning speed of 50 nm/min, with data points taken at 0.1 nm intervals from 190 to 260 nm. Each scan was performed three times and averaged. The spectra of Cyanidin and buffer were subtracted simultaneously. Data visualization, including smoothing and noise reduction, was carried out using Jascow32 software.

### Isothermal titration calorimetry

Isothermal titration calorimetry (ITC) was used to probe the thermodynamic parameters of Peo binding to α-syn at 37 ^o^C, using VP-ITC system (MicroCal, Northampton, MA). The monomeric α-syn (140 μM) sample was centrifuged at 14000 g for 30 min to remove higher order aggregates and the supernatant was degassed for 10 min to remove any air bubbles. Degassed α-syn (140 μM) was added to the ITC cell and continuously titrated with 10 μl of Peo (1500 μM) with a spacing time of 480 s and stirring speed of 524 rpm. Control experiments were performed by titrating Peo with buffer under identical conditions and the heat associated with the reaction was subtracted from α-syn-Peo reaction. The percentage of DMSO (2%) was constant in both cell and syringe in every reaction. High percentage of DMSO was required to solubilise Peo (1500 μM) in the sodium phosphate buffer, pH 7.4. The fitting of the data and baseline corrections were executed using Microcal ORIGIN 7.0 software.

### Size-exclusion chromatography

Samples were prepared by incubating α-syn in the absence and presence of increasing concentrations of Peo (10 - 200 μM) and the larger aggregates were removed by centrifugation at 14000 g for 30 min. The samples were injected into Superdex 200 10/300 GL column (GE Healthcare) with a void volume of 7.8 mL, that was pre-equilibrated with 20 mM sodium phosphate, 100 mM NaCl, and 0.02% sodium azide buffer using Äkta Explorer FPLC instrument (GE Healthcare). Calibration of the column was done using vitamin B12 (1.3 kDa), equine myoglobin (17 kDa), chicken ovalbumin (44 kDa), bovine γ-globulin (158kDa), thyroglobulin (670 kDa) standards (BioRad).

### Cell viability sssay

Human neuroblastoma cells (SH-SY5Y) were grown in a 1:1 mixture of minimal essential medium (MEM) (Sigma-Aldrich) (40%) and nutrient mixture Ham’s F-12 (Sigma-Aldrich) (40%) supplemented with 18% fetal bovine serum (FBS), 1% l-glutamine (3.6 mM), and 1% penicillin/streptomycin antibiotics and grown in a 5% CO2 atmosphere at 37^°^C. The vitality of SH-SY5Y cells after treatment with control and Cyanidin treated α-synwas determined using an MTT [3-(4,5-dimethylthiazol-2-yl)-2,5-diphenyltetrazolium bromide] dye. Live cells with active metabolism convert MTT to formazan. Briefly, cells (25× 10^4^) per well, along with growth medium were seeded onto 96-well poly-L-lysine-coated plates and allowed to adhere for 24 hours at 37°C, followed by incubation of aggregated samples (final concentration 5 μM) were added to the cells and incubated for an additional 48 hours. MTT (5 mg/ml) was added to each well in the dark, followed by the addition of 100% DMSO after 3 hours to dissolve the formazan crystals. After 10 minutes, absorbance was measured at 560 nm, with background subtraction at 650 nm, using a Varioskan Flash microplate reader. The absorbance correlates with the number of viable cells; thus, a vitality greater than 100% may indicate an increase in cell number compared to the control, as these are non-differentiated neuroblastoma cells. The assay was repeated three times, and statistical analysis was performed using Student’s t-test, with p-values above 0.05 considered insignificant.

### Molecular docking

In this study, molecular docking was carried out using AutoDock Vina to evaluate the potential binding interactions between α-synuclein (α-syn) and Cyanidin (Cyan). During the simulations, α-syn was treated as a rigid macromolecule, whereas the ligand was allowed conformational flexibility. Prior to docking, both protein and ligand structures were refined by adding missing hydrogen atoms, assigning charges, bond orders, and flexible torsions where necessary. For the target protein, water molecules were removed and polar hydrogens were introduced to complete the structure. Docking grids were manually defined to ensure that the protein was properly encompassed within the search space. The three-dimensional structure of Cyanidin was retrieved from ChemSpider (ID: 114193). The ligand was docked against seven representative α-syn conformers, which were extracted from the Protein Ensemble Database (PeDB:9AAC) containing NMR- and PRE-derived ensembles. These conformations were selected using the ProFit server, based on RMSD comparisons with a 19 Å cutoff, where one structure was randomly chosen and compared against the remainder of the ensemble. Grid box dimensions for each docking experiment are provided in Table S2. Visualization and interaction analyses of the resulting complexes were performed using Discovery Studio 4.5 and PyMOL.

## Supporting information

Table S1: Calculated Affinity

## AUTHOR INFORMATION

### Author Contributions

G.V. and R.B. designed the research. G.V. performed the experiments. G.V. and R.B. analysed the data and wrote the manuscript.

### Notes

The authors declare no competing financial interest.

## ACKNOWLEDGEMENT

The authors thank Prof. Peter Lansbury (Harvard Medical School) for providing the human α-syn clone and acknowledge Mr. Saroj K. Jha, and Mr. Manu of AIRF and JNU for technical assistance in AFM and TEM experiments, respectively. G.V. thanks CSIR, Government of India for the research fellowship.

## ABBREVIATIONS

PD: Parkinson’s disease
IDP: Intrinsically disordered protein
NAC: Non-amyloid β-component
LB: Lewy body
α-syn: alpha-synuclein
LBD: Lewy body disease
Cyan: Cyanidin
Del: Delphinidin
Peo: Peonidin
ThT: Thioflavin T
TEM: Transmission electron microscopy
AFM: Atomic force microscopy
ANS: 1-anilino-8-naphthalenesulfonate
CD: Circular dichroism
Phe: Phenyl alanine
Tyr: Tyrosine
ITC: Isothermal titration calorimetry
SDS-PAGE: sodium dodecylsulphate-polyacrylamide gel electrophoresis

## Supplementary Material Description

SDS-PAGE analysis of purified α-syn, Nano-ESI mass spectrum of α-syn, ThT assay of control samples, ThT assay of α-syn in the presence of higher concentrations of Cyanidin and Delphinidin, Cytoxicity assay of all three flavonoids, Seeding experiment of the α-syn fibrillar seeds at different concentrations, Molecular docking of Cyanidin with different conformations of α-syn.

