## Supplementary material for "Suppression and modulation mechanism underlying the flavonoid-induced inhibition of fibrillation of α-synuclein": Table S1: Calculated Affinity

### Supplementary Information

### The Anthocyanidin Peonidin Interferes with an Early Step in the Fibrillation Pathway of α-Synuclein and Modulates it towards Amorphous Aggregates

*Geetika Verma*[1] *and Rajiv Bhat*[1*]

[1]School of Biotechnology, Jawaharlal Nehru University, New Delhi, India 110067

AUTHOR INFORMATION

* Corresponding Author

Rajiv Bhat

Professor

School of Biotechnology, Jawaharlal Nehru University, New Delhi, India 110067

**Figure S1:**

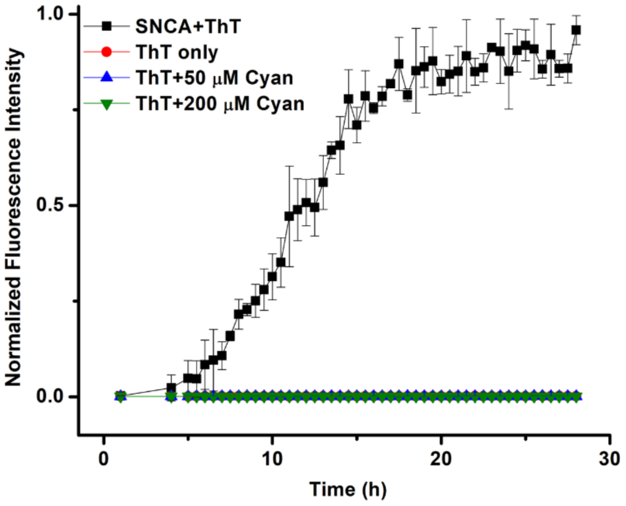


Figure S1: Time-dependent fluorescence intensities measured at 480 nm for ThT alone, ThT with α-syn, ThT with two different concentrations of Cyanidin (50 and 200 μM)

**Figure S2:**

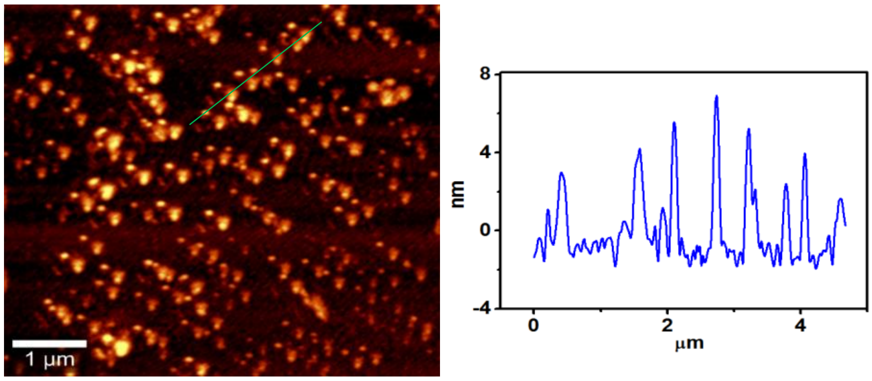


Figure S2: AFM image of On-pathway α-syn oligomers formed during the course of fibrillation.

**Figure S3:**

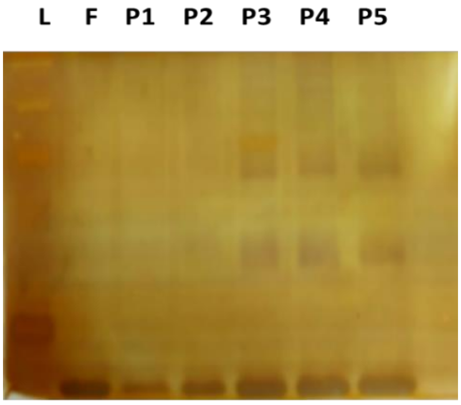


**Figure S3: SDS Polyacrylamide gel electrophoresi**s of the insoluble fraction of the Cyanidin (50 μM) treated α-syn samples, where P1, P2, P3, P4 and P5 correspond to the samples where Cyanidin was added to 0, 3, 12, 24 and 48 hours, respectively, demonstrates the disaggregation of the amyloid fibrils. The molecular weight marker lane in the gel is depicted as L, and F denotes the control α-syn fibrils without Cyanidin.

**Figure S4:**

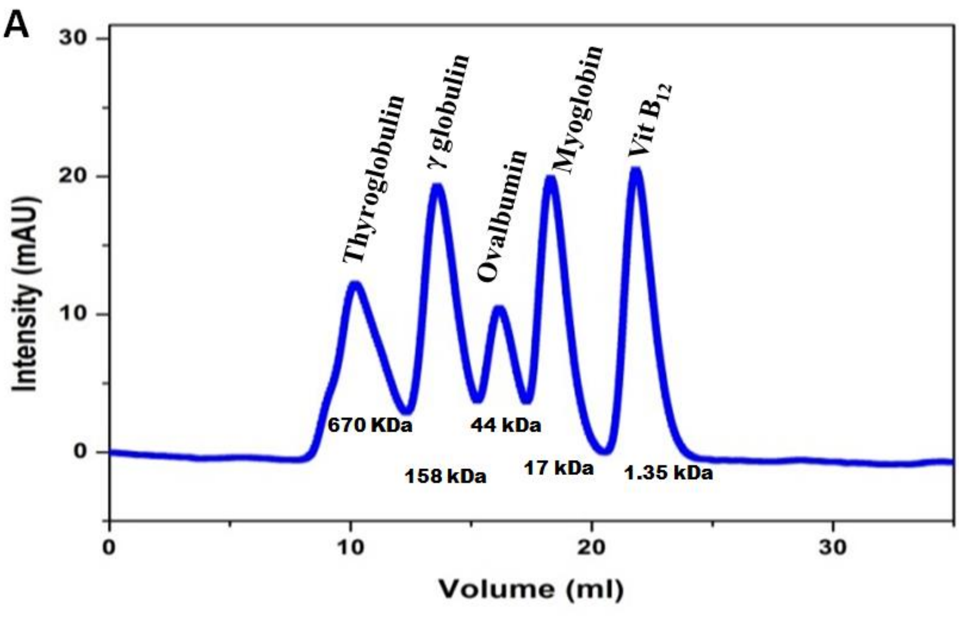


**Figure S4: Size exclusion chromatography. (A)** Elution profile of standard molecular weight protein markers on a superdex 200 10/300 GL size exclusion column.

**Table S1: Calculated Affinity of Hydrogen Bonds (HBs) and Hydrophobic Contacts (HCs) for Specific Regions of the seven conformations of α-synuclein.**

| Interaction | Affinity (kcal/mol) | HBs | Amino Acid | HCs | Amino Acid |
|---|---|---|---|---|---|
| **Conf1-Cyanidin** | **-7.3** | **2** | **Thr 64<br>Gly 68** | **5** | **Lys 60<br>Val 71<br>Ala 76<br>Gln 79<br>Thr 81** |
| **Conf2-Cyanidin** | **-7.3** | **2** | **Lys 32<br>Lys 34** | **3** | **Lys 21<br>Thr 22<br>Tyr 39** |
| **Conf3-Cyanidin** | **-6.0** | **1** | **Glu 110** | **6** | **Val 77<br>Gly 106<br>Gln 109<br>Ile 112<br>Pro 117<br>Asp 119** |
| **Conf4-Cyanidin** | **-7.2** | **1** | **Lys 96** | **10** | **Ala 30<br>Gly 31<br>Lys 32<br>Thr 33<br>Lys 80<br>Ile 88<br>Ala 89<br>Gly 93<br>Ala 90<br>Phe 94** |
| **Conf5-Cyanidin** | **-6.2** | **2** | **Thr 81<br>Lys 96** | **4** | **Lys 80<br>Val 82<br>Asp 98<br>Ile 112** |
